# Liquid-liquid phase separation driven compartmentalization of reactive nucleoplasm

**DOI:** 10.1101/2020.07.13.200980

**Authors:** Rabia Laghmach, Davit A Potoyan

## Abstract

The nucleus of eukaryotic cells harbors active and out of equilibrium environments conducive to diverse gene regulatory processes. On a molecular scale, gene regulatory processes take place within hierarchically compartmentalized sub-nuclear bodies. While the impact of nuclear structure on gene regulation is widely appreciated, it has remained much less clear whether and how gene regulation is impacting nuclear order itself. Recently, the liquid-liquid phase separation emerged as a fundamental mechanism driving the formation of biomolecular condensates, including membrane-less organelles, chromatin territories, and transcriptional domains. The transience and environmental sensitivity of biomolecular condensation are strongly suggestive of kinetic gene-regulatory control of phase separation. To better understand kinetic aspects controlling biomolecular phase-separation, we have constructed a minimalist model of the reactive nucleoplasm. The model is based on the Cahn-Hilliard formulation of ternary protein-RNA-nucleoplasm components coupled to non-equilibrium and spatially dependent gene expression. We find a broad range of kinetic regimes through an extensive set of simulations where the interplay of phase separation and reactive timescales can generate heterogeneous multi-modal gene expression patterns. Furthermore, the significance of this finding is that heterogeneity of gene expression is linked directly with the heterogeneity of length-scales in phase-separated condensates.

## 1. Introduction

Phase separation is a fundamental mechanism for the emergent order in an ordinary and biological matter [1, 2]. Recently, phase separation of bio-molecules has also become a cornerstone physical mechanism for understanding the intracellular organization [2, 3, 4]. A wide range of membrane-less compartments are found to form through biomolecular phase separation, including nucleoli [5, 6, 7], stress granules [8, 9, 10], chromatin domains [11, 12, 13] and transcriptional centers [14, 15]. The primary components driving the intra-cellular phase separation are proteins and nucleic acids with multivalent interaction centers [16, 17].

The stickers-and-spacers framework has emerged as a viable model explaining the existence of a broad class of sequence encoded driving forces of disordered proteins which serve as nucleating centers for biomolecular condensates [3, 18]. These newly appreciated abilities of proteins and nucleic acids for forming large-scale liquid bodies is offering fresh avenues for understanding mechanisms of the coordinated action of biomolecules in gene regulation and cellular organization that go beyond single-molecule action. It is well known that eukaryotic nuclei are rich in disordered proteins linked with transcription and chromatin architecture reorganization activities [19, 20]. There have been several proposals of the functional roles that may include catalysis of biochemical reactions, noise buffering and inducing ultra-sensitive signals [21, 22, 23]. However, understanding the mechanistic picture that links phase separation to the functional gene regulatory processes inside the nucleus has remained elusive [22] due to heterogeneous, multi-scale, and non-equilibrium nature of the nuclear environment [24].

In this work, we propose a minimal model of a reactive nucleoplasm (Fig. 1) with an objective to illustrate, in a proof of principle manner; (i) How spatially resolved non-equilibrium reactive en-events generate qualitatively distinct from equilibrium phase behavior and (ii) How the interplay of various kinetic timescales in the system impacts gene expression patterns. The minimal reactive nucleoplasm model consists of a ternary solution filled with incompressible fluid consisting of “active” protein, RNA components, and “passive” nucleoplasmic buffer. The mathematical formulation of the model is based on a generic ternary diffuse interface model of one-step transcription/translation and phase-separation inside the nucleoplasm. The model resolves the formation and dissolution of protein-RNA droplets as well as reactive events of influx/creation and outflux/degradation of proteins and RNA. By an extensive set of simulations exploring the interplay of timescales in the system, we found a broad kinetic regime dominated by length-scale heterogeneity of phase-separated droplets which correspond to heterogeneous gene expression patterns.

**Figure 1.**
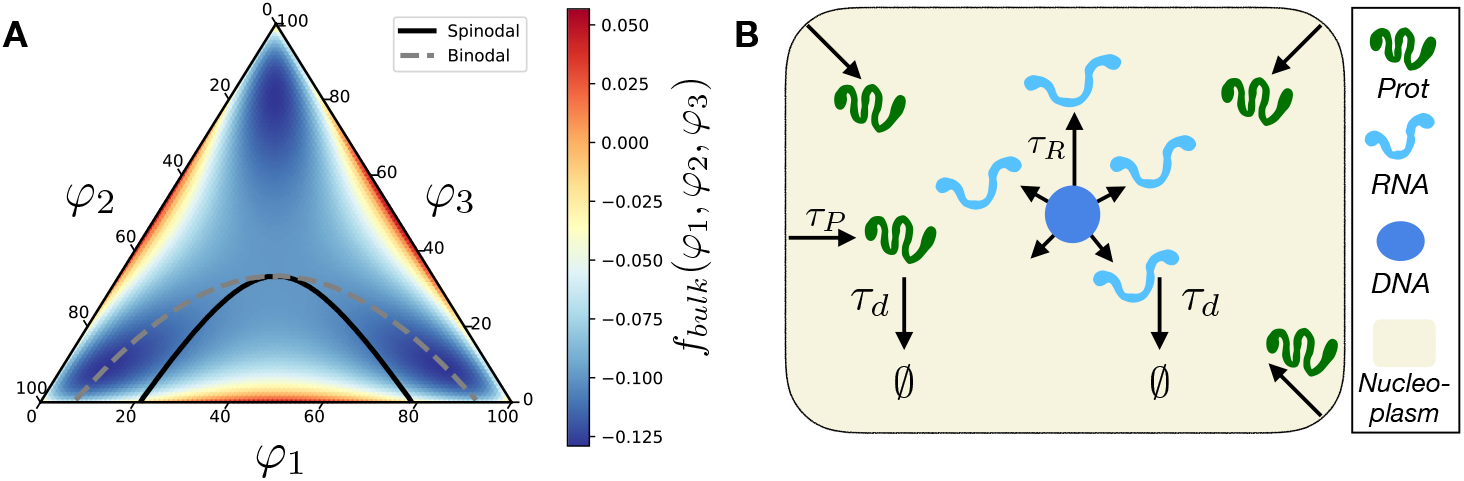
A) Phase diagram of the symmetric ternary protein-RNA-nucleoplasm mixture and the bulk free energy governing the solution thermodynamics *f_bulk_*(*φ*_1_, *φ*_2_, *φ*_3_) for *χ* = 3. The solid-line and dash-line correspond to the spinodal and binodal curves, respectively. B) The schematic of the minimal reactive nucleoplasm model. Shown are the main reactive components (protein, RNA, DNA-seed, nucleoplasm reservoir), the cooresponding reactive processes involving each components as well as their spatial generation profiles

Various simple mathematical models have been used to explore generic aspects of equilibrium and non-equilibrium phase-separation as well as the physical properties of resulting condensates [25, 26, 27, 28, 29, 30, 31, 32, 33]. The most relevant to the present contribution are the work by Berry et al. [29], Tang et al. [33], and Glotzer et al. [32], where authors have considered a binary and ternary fluid model of nucleoplasm, which couples phase separation with first-order exchange among soluble and insoluble components. Authors have simulated different stages of spinodal decomposition and explored its impact on Ostwald ripening of droplets. It was shown that kinetics of the RNA flux accelerates the ripening of droplets, thereby showing a link between the thermodynamics of phase separation and kinetics of droplet formation. Other recent notable studies worth highlighting here include the work by Yamamoto et al. [30], which have studied the non-spatial model of architectural RNA phase separation coupled with non-equilibrium production. Finally, an important theoretical framework by Ilker and Joanny [34] establishes and equivalence of phase-separation kinetics with the Cahn-Hilliard effective temperature.

Some of the key distinctions of the present model from previous studies include (i) Spatially resolved formulation of gene expression and phase separation (ii) Explicit connection of phase-separated patterns with transcription and translation models of gene expression (iii) Exploring the impact of dynamic turnover in RNA-protein-nucleoplasm ternary diagram on global patterning. Thus, the present work clarifies as a first step the dual nature of nuclear order and gene expression, and provides a useful theoretical framework for understanding equilibrium and non-equilibrium origins of intra-nuclear patterning.

## 2. Minimal reactive nucleoplasm model

In order to explore the kinetics of phase separation of RNA-proteins-nucleoplasm reacting mixture, we use a local thermodynamic approach that describes the phase separation of a ternary mixture based on the Flory-Huggins model [35, 36] for polymer solutions coupled with chemical reaction-diffusion equations (Fig. 1). In this section, we describe the mathematical formulation of the model of a “minimal reactive nucleoplasm:” where the liquid-liquid phase separation of single RNA and protein components is coupled with reactive events of transcription, translation, and degradation.

A minimal one-step model of independent, unregulated transcription and translation [37] is used to set the lifetimes of protein and RNA components comprising the nucleopalsmic milieu defined by the following the chemical reactions:

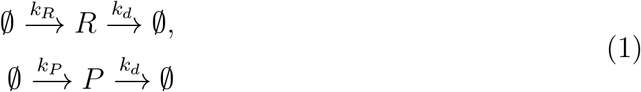

where the *k_R_, k_P_* and *k_d_* are the rate coefficients for transcription, translation and degradation, respectively; The 0 symbol is used to denote the degradation of RNA and protein. The kinetics of phase separation of ternary reactive protein-RNA-nucleoplasm mixture is given by the following set of reaction-diffusion equations [32]:

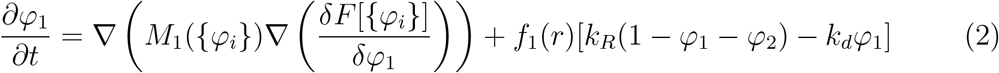

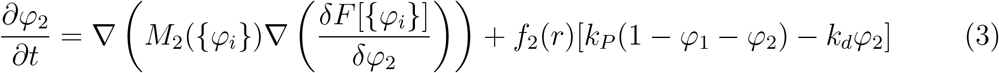

where *φ_i_* (*i* = 1,2,3) are the order parameter variables associated with the local concentration of *i*th-component forming the mixture, *M_i_* are their corresponding mobility coefficients, and *F* is the free energy functional describing the thermodynamics of ternary mixture. The total local density in the system is considered constant, respecting the incompressibility condition expressed as: 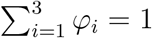. The indices 1, 2, and 3 denote RNA, protein, and nucleoplasm, respectively. The function *f*_1_(*r*) and *f*_2_(*r*) are used to define the spatially heterogeneous distribution of components mimicking RNA expression inside nucleus and protein translation and flow into nucleus. The RNA is created in the center of nucleoplasm thereby mimicking a transcriptional process [38], herein we used the function defined as *f*_1_(*r*) = *exp*(−(*r* – *r_c_*)^2^/*a*) where *r* is the distance from the center of the domain located at *r_c_*. The coefficient a is a localization lengthscale which is set to a=1. The protein component is flown into the nucleus from the nuclear boundaries thereby mimicking translation [38], herein we used the function defined as *f*_2_(*r*) = 1 on the domain boundary *∂*Ω and 0 otherwise. Heterogeneous mobility is an interesting aspect of nucleoplasm [39], however, and is undoubtedly deserving of a separate investigation. For simplicity, we assume the mobility coefficients for all components to be constant *M*_1_({*φ_i_*}) = *M*_2_({*φ_i_*}) = *M*.

The free energy functional taken in this model is based on the Flory-Huggins energy formulation of the ternary mixture system [40, 41], which is given by:

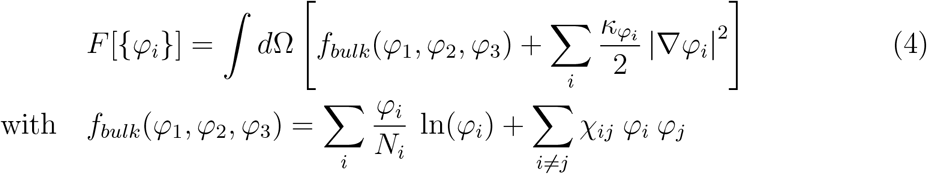

where *f_bulk_*(*φ*_1_, *φ*_2_, *φ*_3_) is the local free energy density, and the square-gradient coefficients *κ_φ_i__* are positive constants controlling the interfacial free energies. *χ_ij_* are the interaction parameters between species *i* – *j*, and *N_i_* is the degree of polymerization. We note here that there also exist other mathematical formulations of the local free energy for describing the phase separation of ternary polymeric mixtures. For instance, the generalized multiwell potential used by Yang et al. [42] to study multiphase systems where the local free energy density is the summation of all the double-well potential for each phase-field variable complemented by a polynomial term that connects all the variables altogether.

The dynamic equations are brought to a dimensionless form, which reveals the essential time- and length-scales of the problem. Let us introduce *l*, the characteristic length of the system, and *τ* is the characteristic time. The dimensionless form of the kinetic equations yields four dimensionless parameters that control the dynamics of phase-separation where components are undergoing chemical reactions. The *τ_D_*/*τ* = *l*^2^/(*M* × *τ*) = 1 is the ratio between diffusion timescale and characteristic time which is fixed to be a unit. The *τ*/*τ_T_* = *τ* × *k_T_* is the ratio between characteristic time and time-scale associated with transcription. The *τ*/*τ_P_* = *τ* × *k_P_* is the ratio between characteristic time and time-scale associated with protein formation. The *τ*/*τ_d_* = *τ* × *k_d_* is ratio between characteristic time and time-scale associated with degradation. To solve the dynamic equations (2)–(3) in the dimensionless form, we use a fully implicit finite element C++ library from the Multiphysics Object-Oriented Simulation Environment (MOOSE) [43]. The simulations were performed on a rectangular domain of dimensions *L_x_* × *L_y_*: (50 × 50), with periodic boundary conditions. A quadrilateral element QUAD4 with four nodes was used for domain meshing with the refinement. The total number of elements used for the fine mesh is 10000. The time step of integration Δ*t* is fixed at 0.05 [a.u.]. In this work, we only consider the case of a symmetric ternary mixture undergoing a chemical reaction with *χ*_12_ = *χ*_23_ = *χ*_13_ = 3 that ensures phase-separation of protein-RNA droplets in the absence of chemical reactions *χ* > *χ_c_*, where *χ_c_* denote the critical point. We also assumed that the chemical reaction takes place at the comparable scales with the phase separation process. Near the critical point *χ_c_*, the mean-field Flory-Huggins free energy can be approximated through Taylor expansion via Landau form [32]. It is important to construct representative free energy functional that accounts for fluctuations of the phase-field variables that are dominant near the critical point. To express the critical behavior a new formulation of the free energy of mixing renormalized by the spatial variation has been proposed by Yamamoto et al. [44, 45]. The chain lengths *N_i_*, or degree of polymerization, are assumed to be the same: *N*_1_ = *N*_2_ = *N*_3_ = 1. It is noted here that the entropic term of the local free energy is proportional to the inverse polymeric lengths *N_i_*. As a consequence of the reduction of entropy for long-chain lengths, the phase diagrams and the spinodal curve will be modified in the way to increase the separating-phase region of the phase diagram. In this case, a small variation of the interaction parameter between species will lead easily to phase separation. The impact of phase-separation of small and large chains such as genomic regions and proteins/RNA could be an interesting study to address in future investigations. The other parameters of simulations were set to *κ*_*ϕ*_1__ = *κ*_*ϕ*_2__ = 0.03125. The initial configuration of phase-field variables is generated by *φ_i_*(*r*) = 〈*φ_i_*〉 + *δφ_i_*, where 〈*φ_i_*〉 is an initial average concentration associated with *φ_i_*, and *δφ_i_* is a small random perturbation amplitude. For all simulations presented here, the initial average of species *i* are set to 〈*φ*_1_〉 = 0.3 and 〈*φ*_2_〉 = 0.09, with *δφ*_1_ ∈ [−0.05, 0.05] and *δφ*_2_ ∈ [−0.01, 0.01].

## 3. Results

Here we report the findings obtained by analyzing the results of simulations with a minimal reactive nucleoplasm model. We have organized the results in three broad kinetic regimes, which are described by three orders of magnitude in degradation time-scale *τ_d_* = *τ*, 10*τ*, 100*τ* with respect to the diffusion time-scale *τ*. These time-scales correspond to different rates of turnover of nucleoplasmic components from fast to slow. For each regime, we have carried out extensive grid-based kinetic parameter sweeps exploring the coupling of transcriptional and transnational time-scales on the backdrop of fixed phase-separating free energy landscape of RNA-protein-nucleoplasm components. Despite the incredible simplicity of the minimal reactive nucleoplasm model, the time course of simulations (Fig. 2–4) has revealed a non-trivial patterning of nucleoplasm which are a dramatic departure from equilibrium thermodynamics of ternary phase separation in the absence of spatially non-uniform reaction-diffusion.

**Figure 2.**
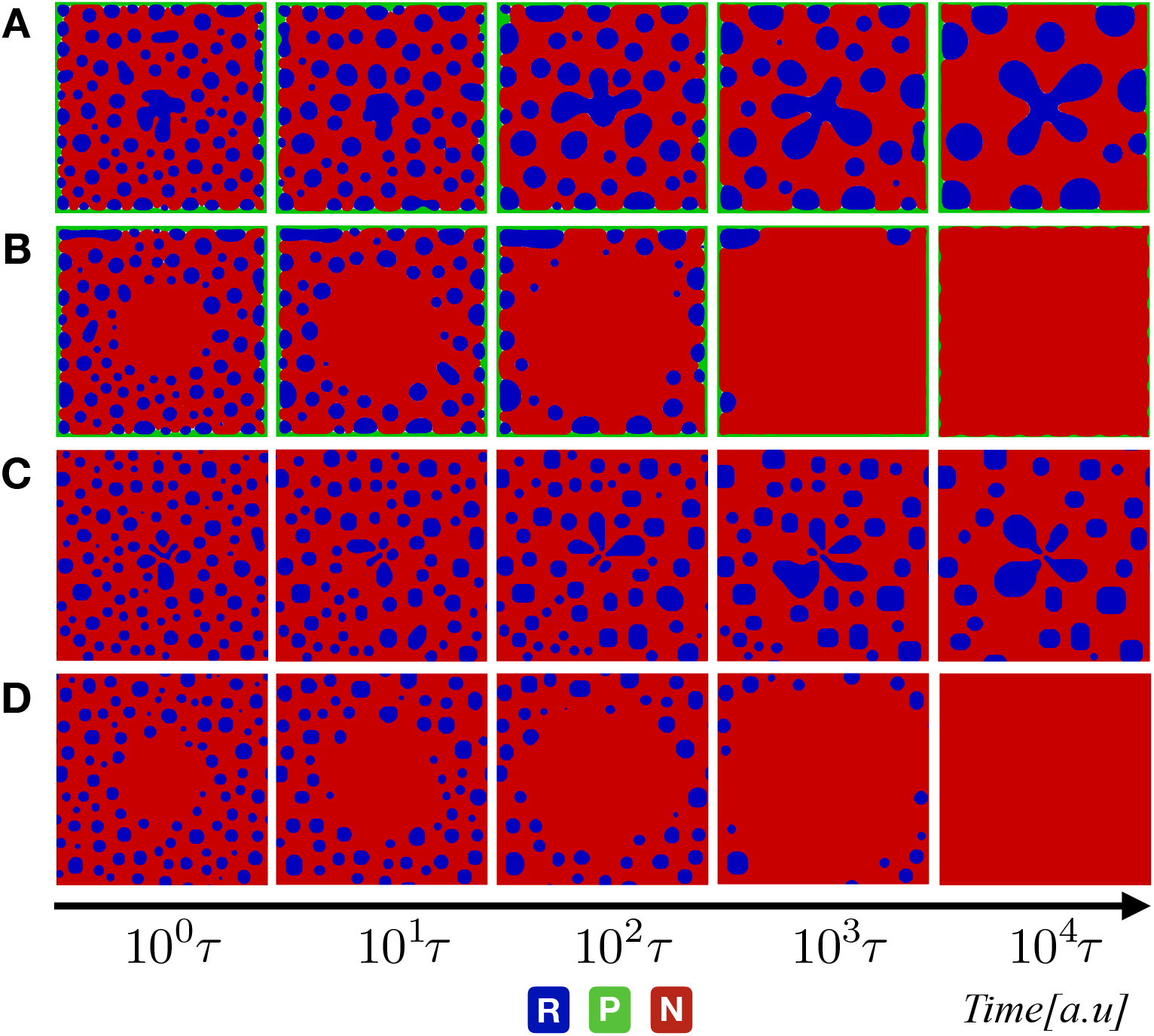
Evolution of phase-field variables *φ_i_* for three phase mixture undergoing the chemical reaction with a degradation time-scale fixed at the same of diffusion (*τ_d_* = *τ*). From up to bottom: snapshots corresponding to simulation results with A) *τ_R_* = *τ_P_* = *τ*; B) *τ_R_* = 100*τ_P_* = 100*τ*; C) *τ_P_* = 100*τ_R_* = 100*τ*; and D) *τ_R_* = *τ_P_* = 100*τ*. The color code in blue, green, and red indicates the RNA, protein, and nucleoplasm regions, respectively.

We start by investigating the fast dynamical regime corresponding to *τ_d_* = *τ*. Fig. 2 shows the spatial and temporal evolution of RNA-protein components in a reactive nucleoplasm environment. Four interesting cases are highlighted in Fig. 2, corresponding to different time-scale separation between translation *τ_P_* and transcription *τ_R_* processes. When *τ_P_* = *τ_R_* = *τ* we observe rapid coarsening dynamics (droplet growth kinetics) and emergence of RNA droplets with stable “protein front” by which we refer to the formation of protein layer surrounding nucleoplasm domain edges (Fig. 2A). Coarsening dynamics for either protein front or RNA droplets are also observed for fast translation/slow transcription or slow translation/fast transcription, respectively (Fig. 2BC). Naturally, it is expected that the more rapid degradation timescales for RNA and protein lead to the disappearance of both RNA droplets and protein fonts(Fig. 2D). Thus, we can conclude that in the fast dynamical regime, nucleoplasm patterns are entirely set by the kinetic parameters with thermodynamic free energy landscape taking the back seat.

The formation of RNA-protein patterns is more intricate in the intermediate dynamical regime corresponding to *τ_d_* = 10*τ* (Fig. 3). In the Fig. 3, we show the patterns arising from competing transcription, translation, and degradation with different transcription and translation timescales. When both transcription and translation have comparable time scales (Fig. 3A) to that of diffusion, we find a rapid progression of protein front on the one hand and a rapid RNA seed droplet growth at on the other. In this regime, once the nucleoplasm reaches a non-equilibrium steady state, droplet patterning is now becomes dictated by the thermodynamics of Flory-Huggins interaction parameters, which set the extend of mixing between protein and RNA components. When both the transcription and translation have the same time scale but are now significantly slower than diffusion(Fig. 3D), then the system displays heterogeneity in protein RNA droplet distribution. The situation with faster transcription and the faster translation is predictable: (Fig. 3B and C) the system tends to increase in protein front and RNA nucleating center, respectively. We have only highlighted the most interesting cases for each dynamical regime. The simulation results for the full set of time-scales are presented in the Supporting Information Fig S1-S10. The temporal evolution of average concentration for different kinetic regimes with *τ_R_* = *τ_P_* = 100*τ* is shown in figure S14.

**Figure 3.**
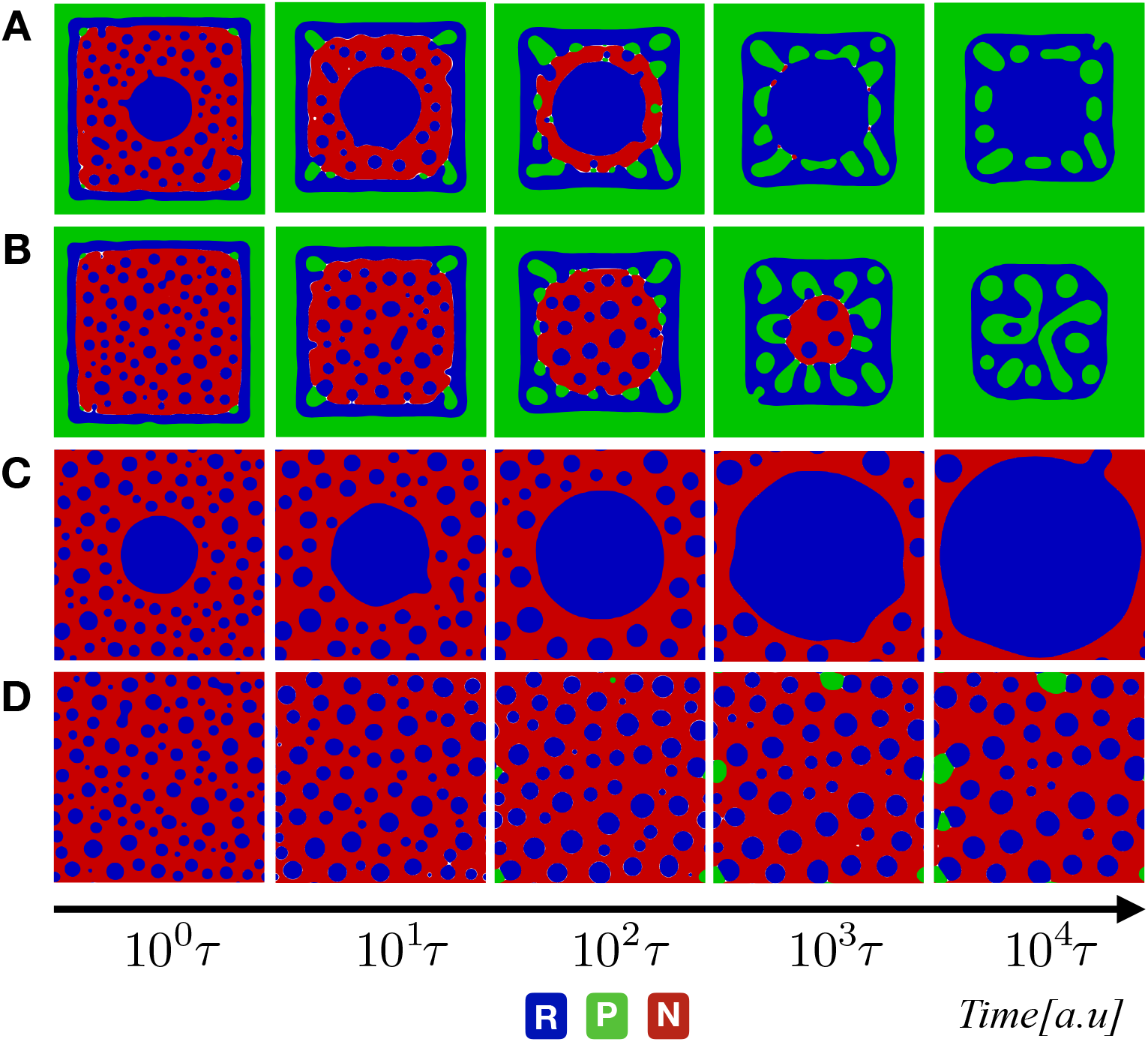
Evolution of phase-field variables *φ_i_* for three phase mixture undergoing the chemical reaction with a degradation time-scale fixed at the same of diffusion (*τ_d_* = 10*τ*). From up to bottom: snapshots corresponding to simulation results with A) *τ_R_* = *τ_P_* = *τ*; B) *τ_R_* = 50*τ_P_* = 50*τ*; C) *τ_P_* = 100*τ_R_* = 100*τ*; and D) *τ_R_* = *τ_P_* = 50*τ*. The color code in blue, green, and red indicates the RNA, protein, and nucleoplasm regions, respectively.

We now turn to the analysis of the slow dynamical regime corresponding to a degradation timescale *τ_d_* = 100*τ*. In this regime, we once again highlight four interesting cases (Fig. 4A-D). When the system has comparable timescales for transcription, translation, and diffusion *τ_P_* = *τ_R_* = *τ*, the nucleoplasmic component gets eliminated, and the system rapidly reaches a balance between protein front and RNA seed. Slowing down the translation *τ_P_* = 100*τ_R_* = 100*τ* by two orders of magnitude leads to a nontrivial patterning with the RNA seed, nucleoplasm and RNA droplets all con-existing at a non-equilibrium steady-state. On the other hand, slowing down the transcription (Fig 4C) leads to a steady-state with significantly downsized RNA-seed at the expense of the peripheral protein front. Slowing down both transcription and translation (Fig 4D) leads to a non-trivial patterning with protein front, RNA droplets, and nucleoplasmic environment, but this time with no dominant RNA-seed.

**Figure 4.**
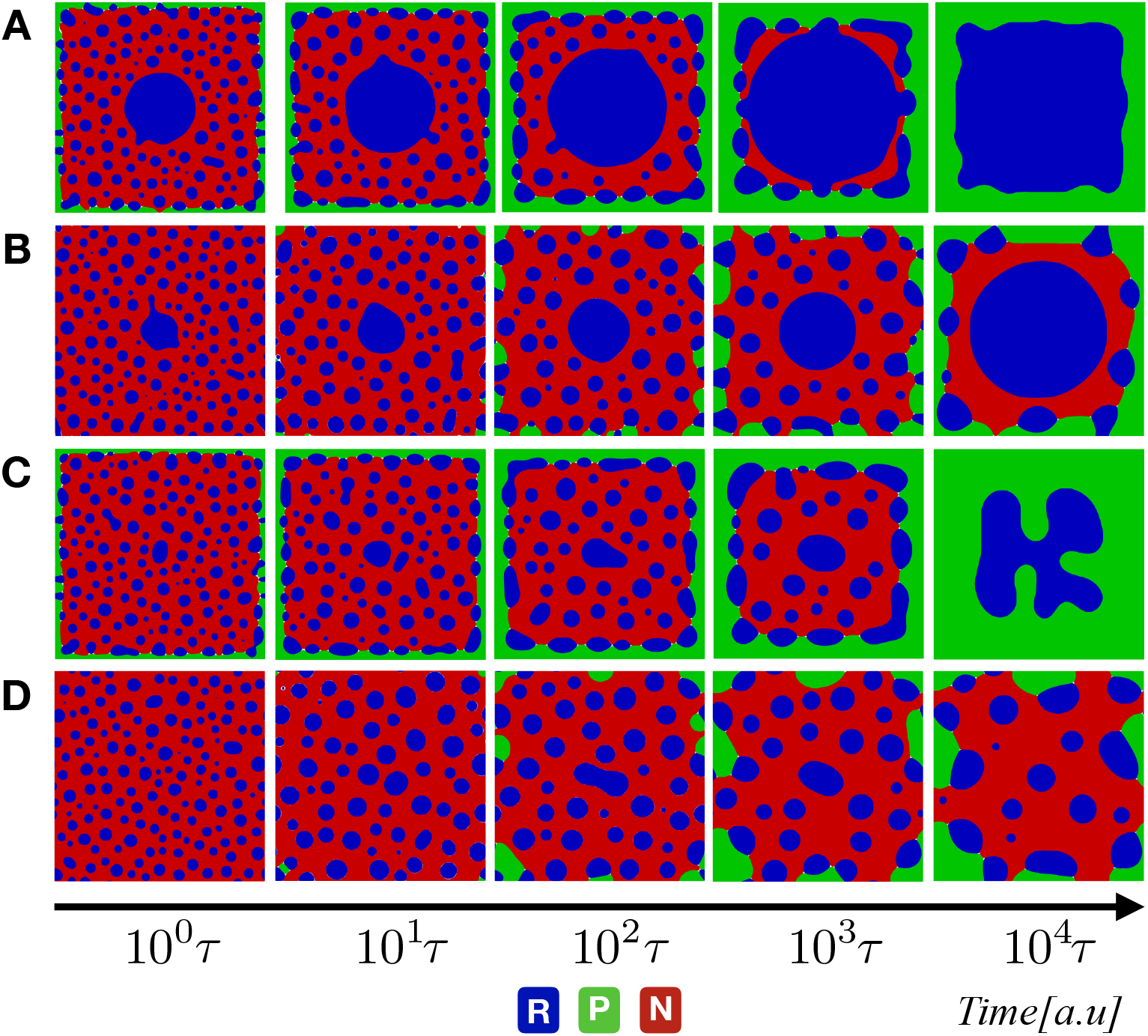
Evolution of phase-field variables *φ_i_* for three phase mixture undergoing the chemical reaction with a degradation time-scale fixed at the same of diffusion (*τ_d_* = 100*τ*). From up to bottom: snapshots corresponding to simulation results with A) *τ_P_* = 10*τ_R_* = 10*τ*; B) *τ_R_* = 10*τ τ_P_* = 50*τ*; C) *τ_R_* = 50*τ τ_P_* = 10*τ*; and D) *τ_R_* = *τ_P_* = 100*τ*. The color code in blue, green, and red indicates the RNA, protein, and nucleoplasm regions, respectively.

To quantify the emergence of different droplet patterns, we have summarized the phase behavior of a reactive nucleoplasm model via. Global kinetic phase diagram, Fig. 5. Here one can see a global picture of how the interplay of kinetic timescales is favoring uniform vs. binary vs. ternary phases.

**Figure 5.**
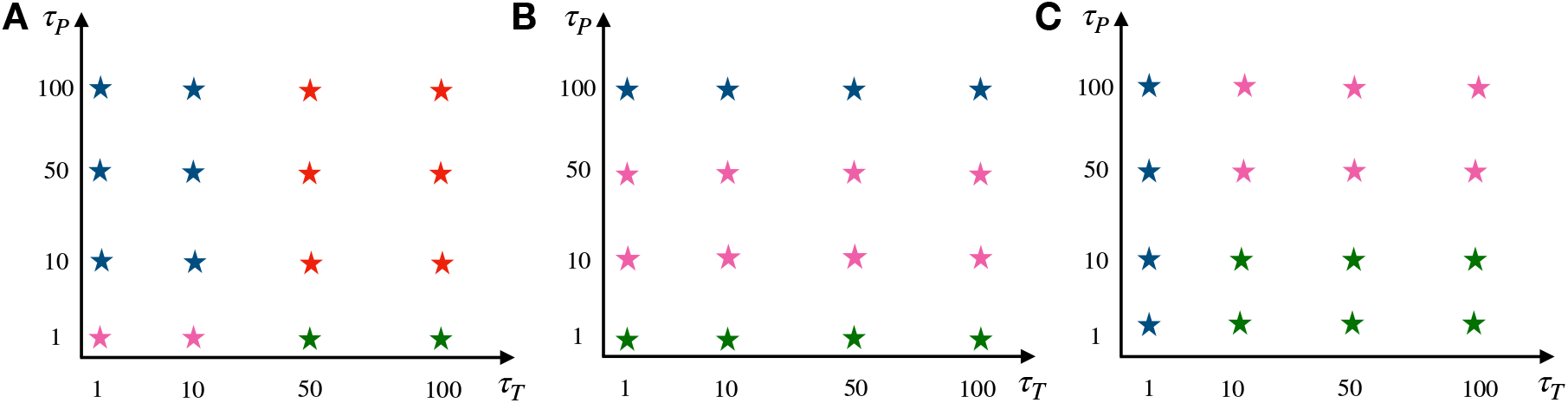
Phase diagram: Phase diagram showing the dominant steady-state phase and summarizing various patterns that arise at three kinetic regimes (A) *τ_d_* = *τ*; B) *τ_d_* = 10*τ*; C) *τ_d_* = 100*τ*). The blue star indicates that the RNA domain becomes the dominant phase in the long timescale limit. The green star means that the proteins-droplets become the dominant phase in the long timescale limit. The pink star indicates three coexisting phases of the ternary mixture. In contrast, the red star indicates that RNA-proteins turn totally into the nucleoplasm.

In order to quantify the length-scales of emergent patterns, we have analyzed the azimuthally averaged dynamic structure factors *S*(*k, t*) = ∪ *dk*_Ω_*S*(**k**, *t*) associated with each dynamical regime (See Fig. 6 and Supporting Information, Fig S11-S13). Analyzing dynamic structure factors reveals characteristic length-scales of protein/RNA droplets as well as emergent dynamical heterogeneity manifesting in the fast or slow coarsening of the droplets. The computed structure factors show a broad range of kinetically controlled states where one has bi-modality of RNA (or protein) components. This bimodality or more broadly heterogeneity of distribution is directly linked to the heterogeneity of droplet sizes and shapes that one can quantify from simulation images (Fig. 3,4). Multi-modal gene expression is a feature often linked with phenotypic heterogeneity and which has been mostly explained by citing the underlying non-linear dynamics of dichotomous switching noise in a spatially uniform master equation formalism [46, 47, 48]. The results of Fig. 6 clearly show that transcriptional heterogeneity can originate purely from the spatially non-uniform nature of gene expression, which is being modulated by timescales of phase separation, transcription, and translation.

**Figure 6.**
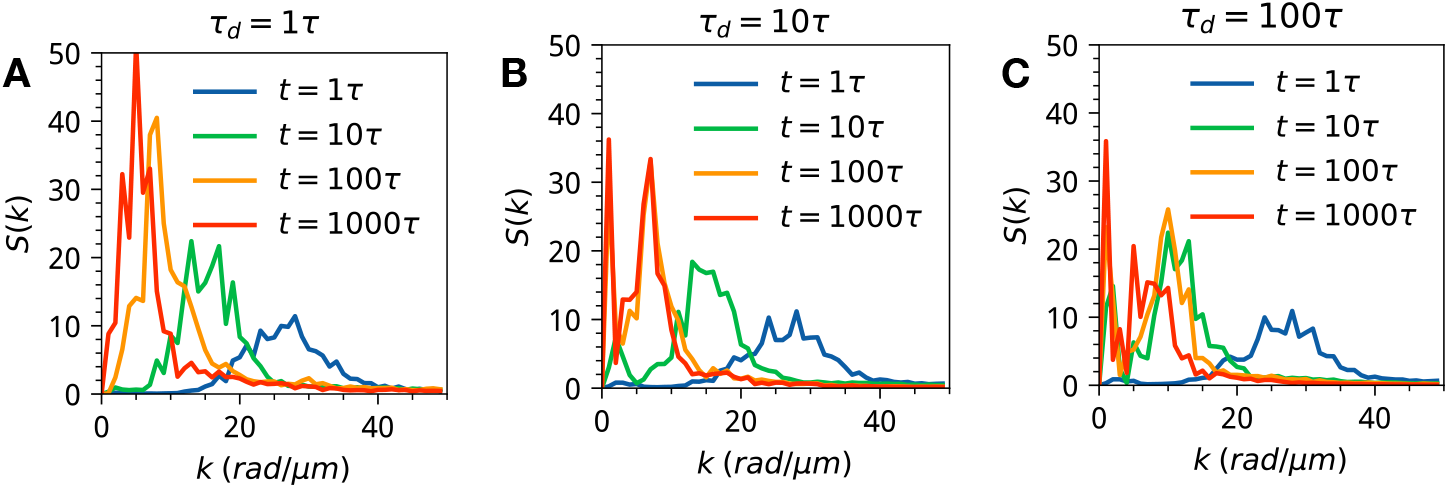
The dynamic structure factors for the three representative kinetic regimes with significant time-scale disparity between transcription and translation *τ_P_* = 100*τ_R_* (See Supporting Information for all the results). The inset shows the scaling and power law exponent. The three panels stand for three dynamical regimes: (A) *τ_d_* = *τ* (B) *τ_d_* = 10*τ* (C) *τ_d_* = 100*τ*.

Finally, we have computed the length scale of patterns summarizing transitional heterogeneity in three kinetic regimes of the minimal reactive nucleoplasm model (Fig. 7). The length-scales patterns are quantified by using pre-computed azimuthally averaged dynamic structure factors 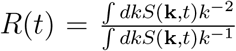 [49]. The length-scale patterns *R*(*t*) quantify the dominant length-scales which emerge during the time-evolution of a reactive nucleoplasm. Analyzing the evolution of length-scale patterns for three dynamical regimes, we see clearly that for the fast dynamical regime, there is only one dominating length-scale, which is dictated by the rapid degradation kinetic timescale. For the intermediate and slow dynamical regimes, however, we find heterogeneity of length-scales, which emerges from the disparity in transcription and translational timescales. We note that this heterogeneity has both structural and dynamical manifestations, as one can see by analyzing the power low of length-scale patterns *R*(*t*) ~ *At^α^*. We find two exponents, one which is characteristic for the early coarsening stage (~ 1/3) and second (*α* ~ 3/8) for the later accelerated phase separating evolution towards a steady state. We note that power laws in the dynamical variables of the nucleoplasmic environment have been detected and characterized in a large number of experiments [39, 50]. These power-law dependencies, however, are hard to disentangle in terms of distinct contributions since their origin may come from any combination of phase-separation, polymeric effects, confinement, and non-equilibrium motorized activities in the nucleus. In this work, we have only managed to scratch at the surface of the fascinating dynamical patterning potential of the active nucleoplasmic environment. For future studies, it would be interesting to investigate the impact of differential mobility, hydro-dynamical coupling, and as well as investigate gene expression beyond a simple one-step model of unregulated reactions that have not been done in the present contribution.

**Figure 7.**
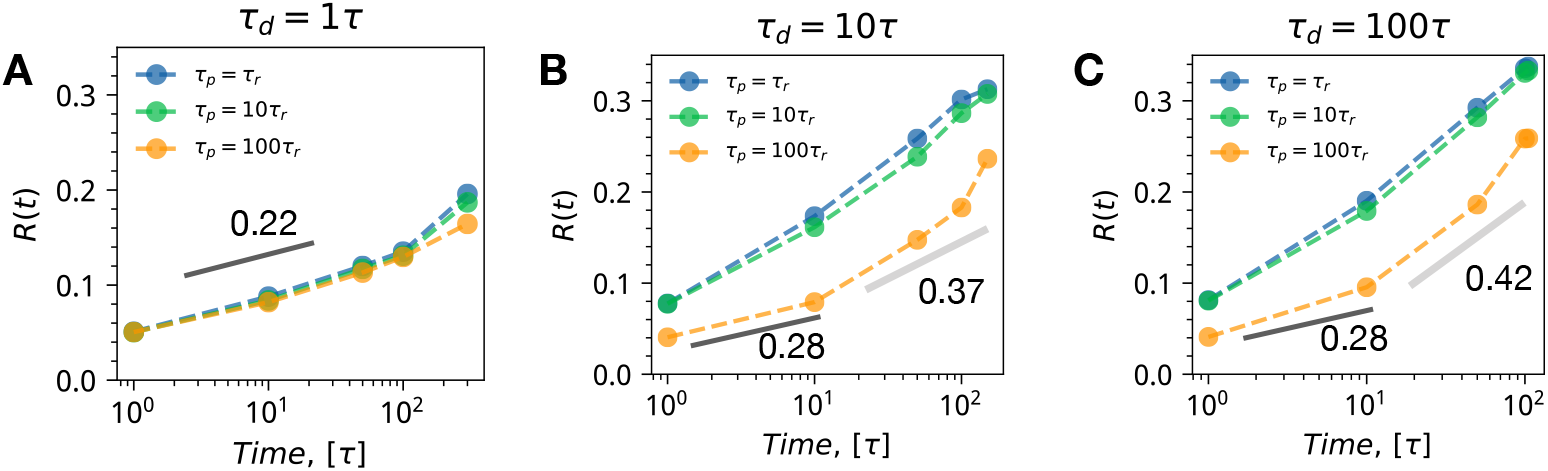
The dominant length-scale patterns for the three representative kinetic regimes. Each panel shows a combination of transcription and translation time-scales sorted with an increasing time-scale disparity from blue to green to orange. (A) *τ_d_* = *τ* (B) *τ_d_* = 10*τ* (C) *τ_d_* = 100*τ*.

## 4. Conclusion

Recent in vitro experiments with binary and ternary protein and RNA mixtures [51, 52, 53] have shown the rich complexity of phases which can emerge through liquid-liquid phase separation via modulation of stoichiometry of components and point mutations. These experiments show potentially novel regulatory strategies which when combined with non-equilibrium cellular processes such as transcription and translation could produce coordinated gene regulatory and signaling actions.

To this end, in the present contribution we introduce a minimal reactive nucleoplasm model combining spatially dependent transcription and translation to liquid-liquid phase separation of RNA and protein components embedded in the backdrop of passive nucleoplasm buffer. We use the minimal reactive-nucleoplasm model to cleanly dissect how the interplay of transcription, translation, and degradation time scales couples with the liquid-liquid phase separation of RNA and protein components in a model ternary solution. By carrying out extensive grid-based sweeps of kinetic parameter space, we uncover various non-trivial patterning and lengthscale heterogeneity compared to the classic Flory-Huggins thermodynamic picture for ternary polymeric solutions. Our central finding is the existence of a broad kinetic regime characterized by a slow turnover of components and timescale disparity between transcription and translation under which a phase separating system can display bi-modal distribution. The significance of this finding is that the observed heterogeneity of gene expression is linked directly with the heterogeneity of length-scales in phase-separated condensates. The main findings and the minimal reactive nucleoplasm model thus establishes a useful framework with which one can further elucidate the emergence of nucleoplasm patterns and phenotype heterogeneity from first principles modeling of phase-separation and reaction-diffusion processes.

## Acknowledgments

Research reported in this publication was supported by the National Institute Of General Medical Sciences of the National Institutes of Health under Award Number R35GM138243. The content is solely the responsibility of the authors and does not necessarily represent the official views of the National Institutes of Health.

This work used the Extreme Science and Engineering Discovery Environment (XSEDE), which is supported by the National Science Foundation grant number ACI-1548562 [54] on the Stampede2 machine at the Texas Advanced Computing Center (TACC) through allocation CTS190023. The authors acknowledge the Texas Advanced Computing Center (TACC) at The University of Texas at Austin for providing HPC resources that have contributed to the research results reported within this paper. The authors also acknowledge financial support from Iowa State University.

## Notes

### Competing Interest Statement

The authors have declared no competing interest.

