## Supplementary Information for "Liquid-liquid phase separation driven compartmentalization of reactive nucleoplasm" (from another version of this preprint: version 1, 2020-07-14)

### Supporting Information for: "Liquid-liquid phase separation driven compartmentalization of reactive nucleoplasm."

July 13, 2020

#### 1 Supporting Information

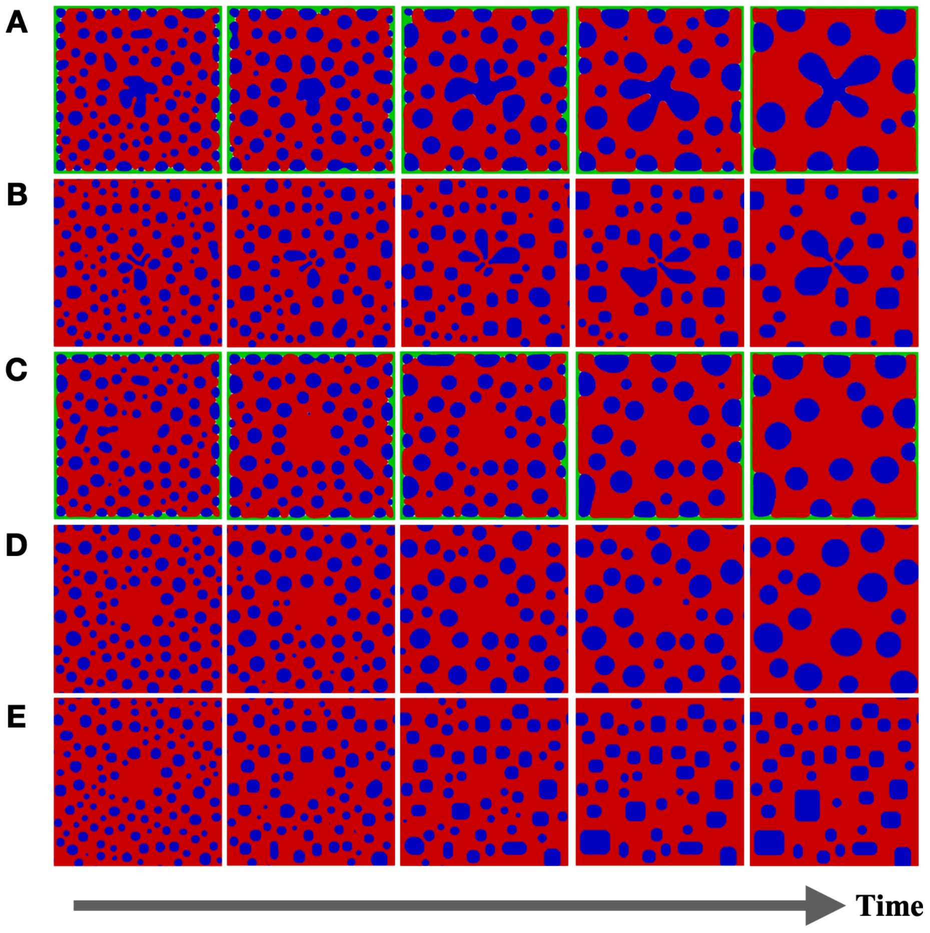


Figure 1: Evolution of phase-field variables $\varphi_i$ for **a fast dynamical regime** $\tau_d = \tau$. From up to bottom: snapshots corresponding to simulation results with A) $\tau_R = \tau_P = \tau$; B) $\tau_P = 100\tau_R = 100\tau$; C) $\tau_R = 10\tau_P = 10\tau$; D) $\tau_R = \tau_P = 10\tau$; and E) $\tau_P = 10\tau_R = 100\tau$.

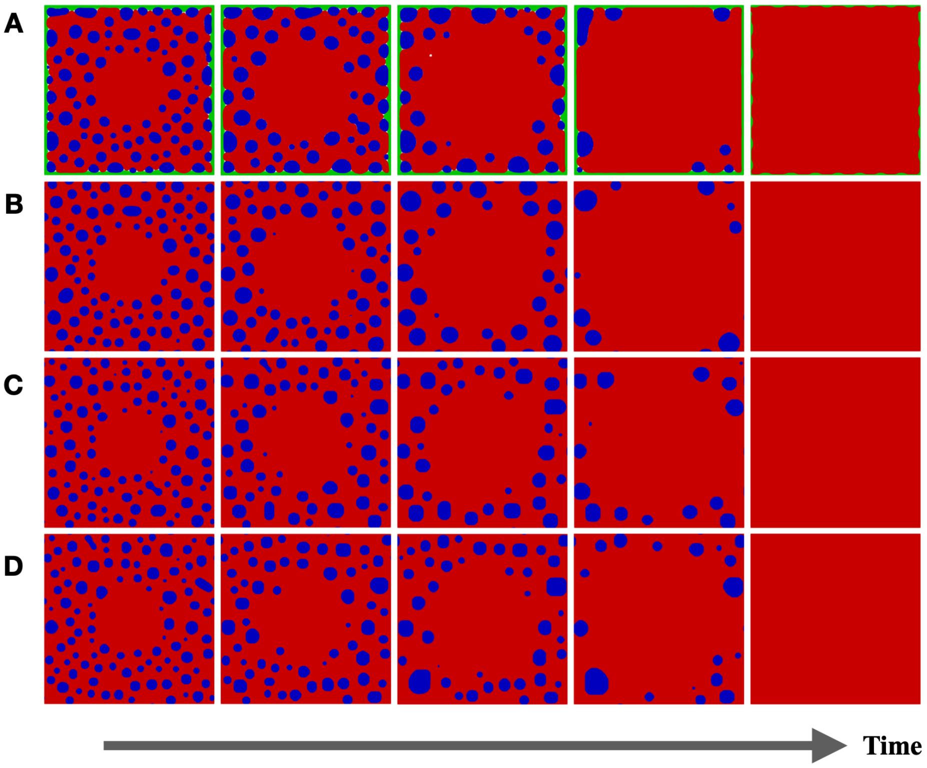


Figure 2: Evolution of phase-field variables $\varphi_i$ for **a fast dynamical regime** $\tau_d = \tau$ **with** $\tau_R = 50\tau$. From up to bottom: snapshots corresponding to simulation results with A) $\tau_P = \tau$; B) $\tau_P = 10\tau$; C) $\tau_P = 50\tau$; and D) $\tau_P = 100\tau$.

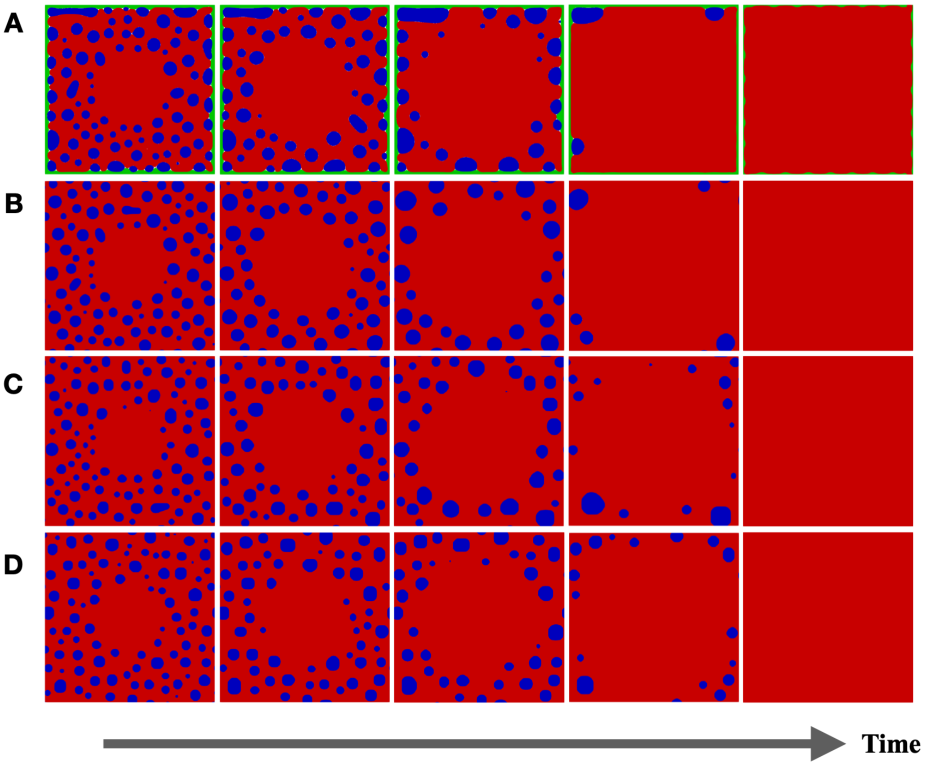


Figure 3: Evolution of phase-field variables $\varphi_i$ for **a fast dynamical regime** $\tau_d = \tau$ **with** $\tau_R = 100\tau$. From up to bottom: snapshots corresponding to simulation results with A) $\tau_P = \tau$; B) $\tau_P = 10\tau$; C) $\tau_P = 50\tau$; and D) $\tau_P = 100\tau$.

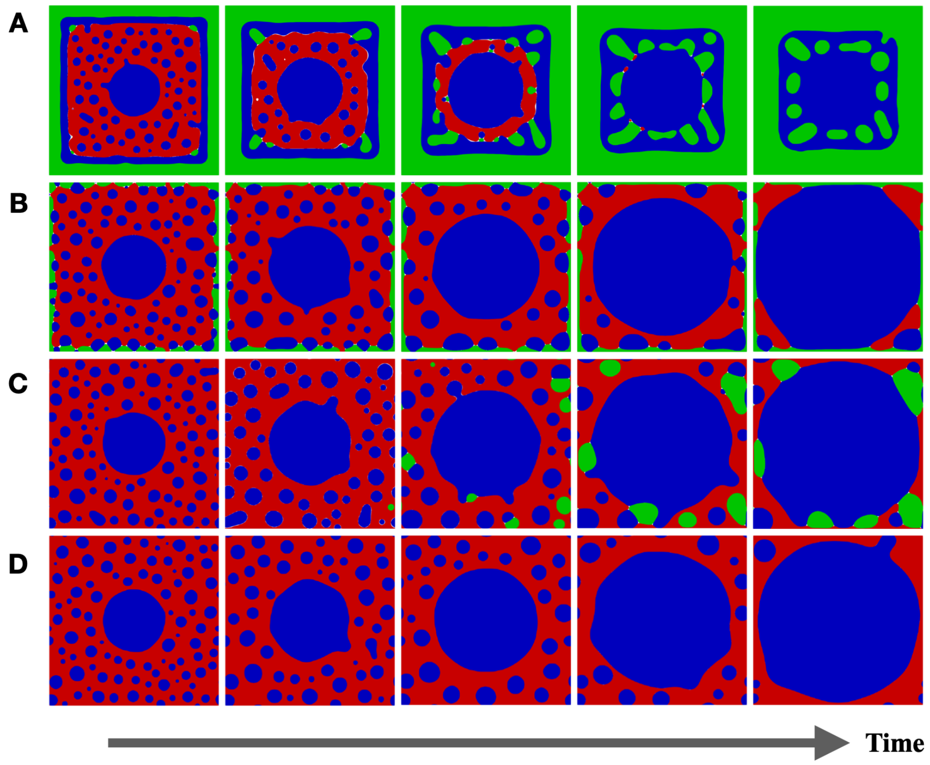


Figure 4: Evolution of phase-field variables $\varphi_i$ for **an intermediate dynamical regime** $\tau_d = 10\tau$ **with** $\tau_R = \tau$. From up to bottom: snapshots corresponding to simulation results with A) $\tau_P = \tau$; B) $\tau_P = 10\tau$; C) $\tau_P = 50\tau$; and D) $\tau_P = 100\tau$.

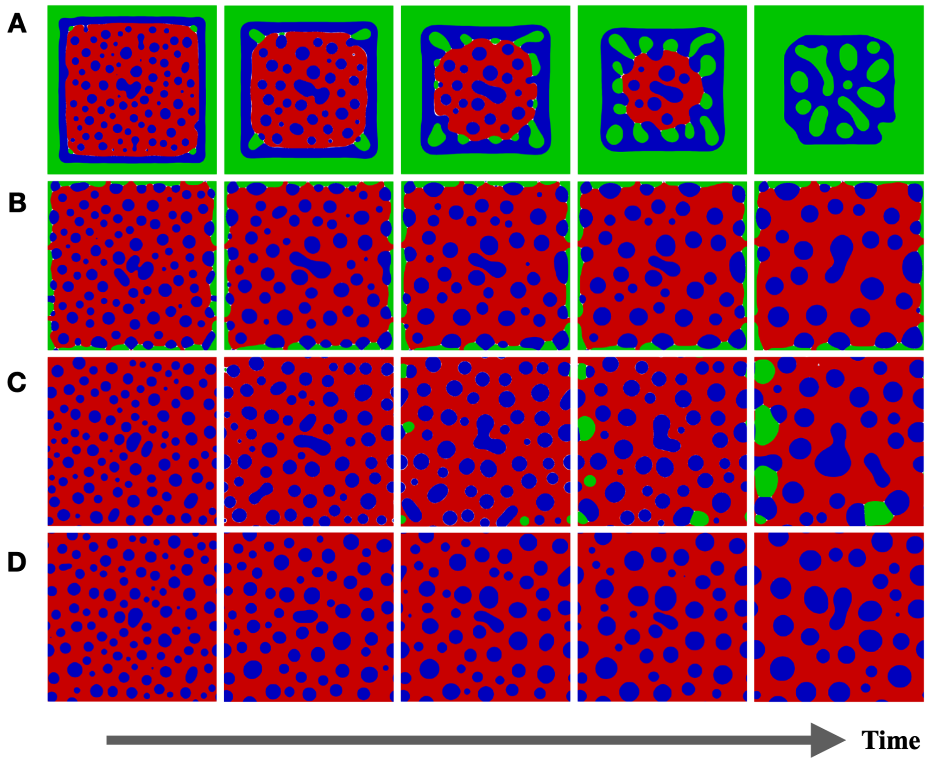


Figure 5: Evolution of phase-field variables $\varphi_i$ for **an intermediate dynamical regime** $\tau_d = 10\tau$ **with** $\tau_R = 10\tau$. From up to bottom: snapshots corresponding to simulation results with A) $\tau_P = \tau$; B) $\tau_P = 10\tau$; C) $\tau_P = 50\tau$; and D) $\tau_P = 100\tau$.

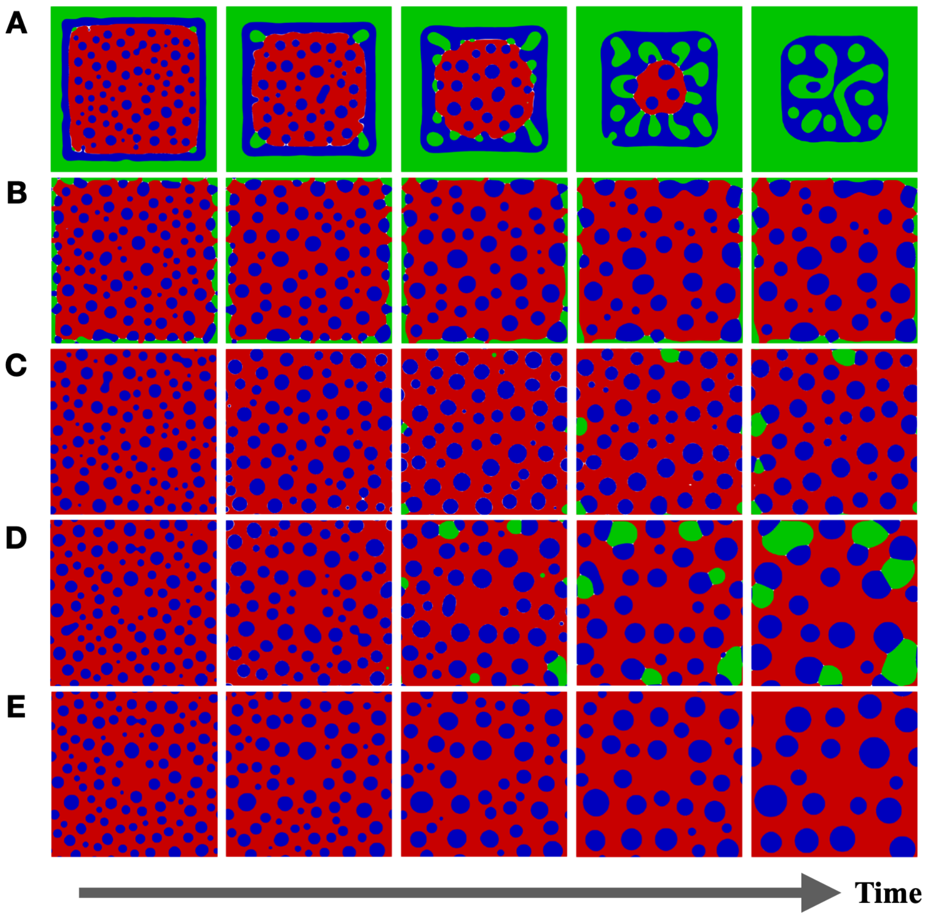


Figure 6: Evolution of phase-field variables $\varphi_i$ for **an intermediate dynamical regime** $\tau_d = 10\tau$. From up to bottom: snapshots corresponding to simulation results with A) $\tau_R = 50\tau_P = 50\tau$; B) $\tau_R = 5\tau_P = 50\tau$; C) $\tau_R = \tau_P = 50\tau$; D) $\tau_R = 2\tau_P = 100\tau$; and E) $\tau_R = \tau_P = 100\tau$.

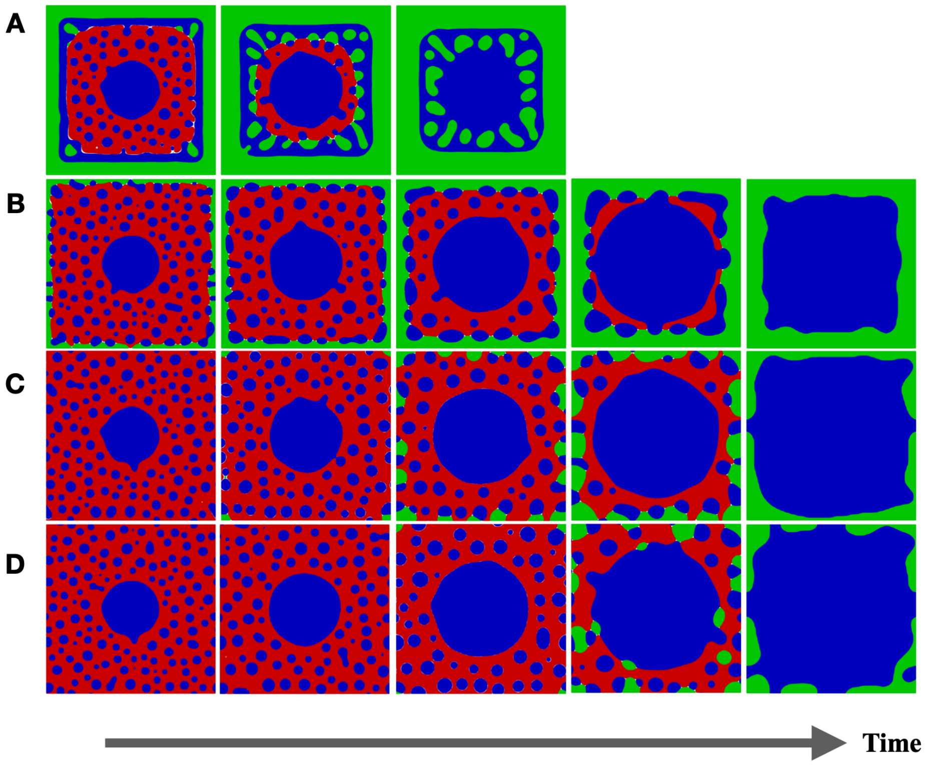


Figure 7: Evolution of phase-field variables $\varphi_i$ for **a slow dynamical regime** $\tau_d = 100\tau$ **with** $\tau_R = \tau$. From up to bottom: snapshots corresponding to simulation results with A) $\tau_P = \tau$; B) $\tau_P = 10\tau$; C) $\tau_P = 50\tau$; and D) $\tau_P = 100\tau$.

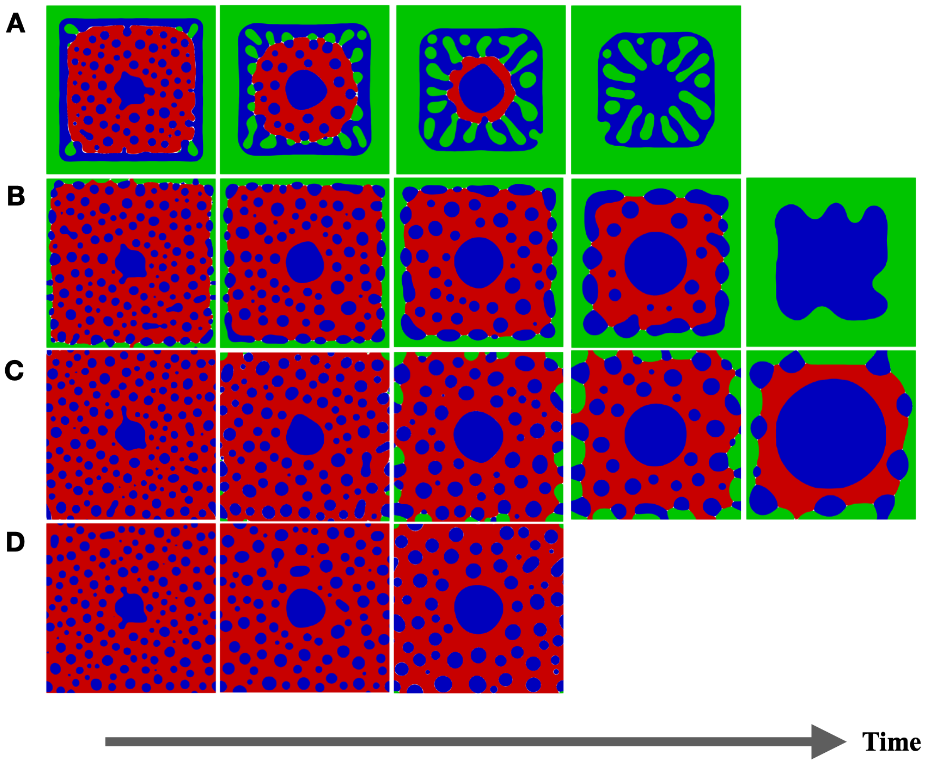


Figure 8: Evolution of phase-field variables $\varphi_i$ for **a slow dynamical regime** $\tau_d = 100\tau$ **with** $\tau_R = 10\tau$. From up to bottom: snapshots corresponding to simulation results with A) $\tau_P = \tau$; B) $\tau_P = 10\tau$; C) $\tau_P = 50\tau$; and D) $\tau_P = 100\tau$.

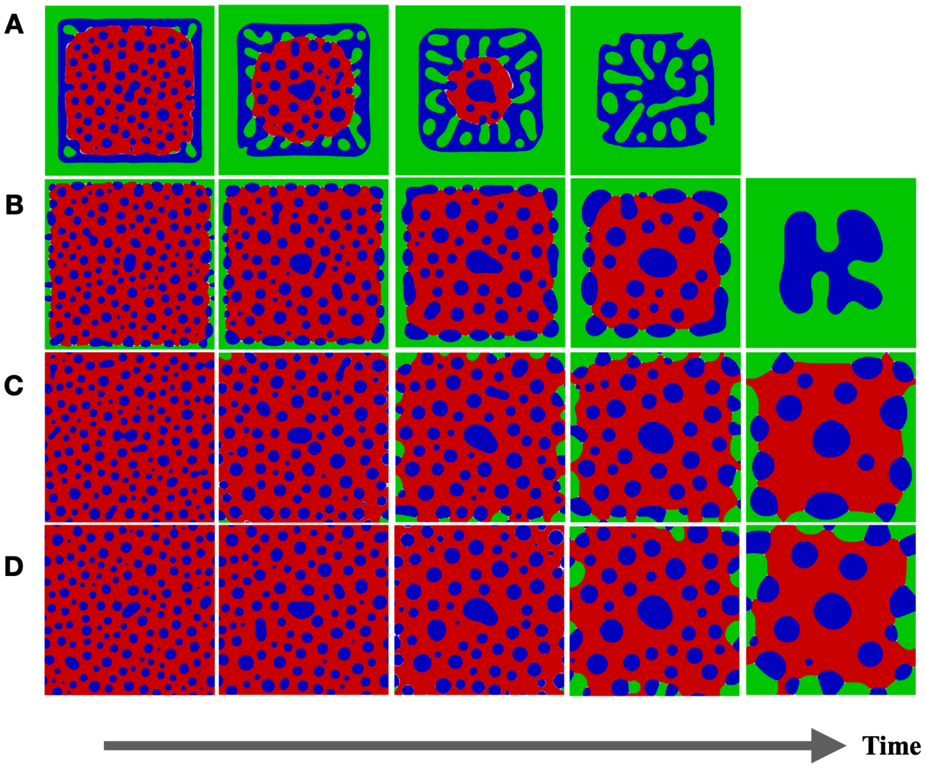


Figure 9: Evolution of phase-field variables $\varphi_i$ for **a slow dynamical regime** $\tau_d = 100\tau$ **with** $\tau_R = 50\tau$. From up to bottom: snapshots corresponding to simulation results with A) $\tau_P = \tau$; B) $\tau_P = 10\tau$; C) $\tau_P = 50\tau$; and D) $\tau_P = 100\tau$.

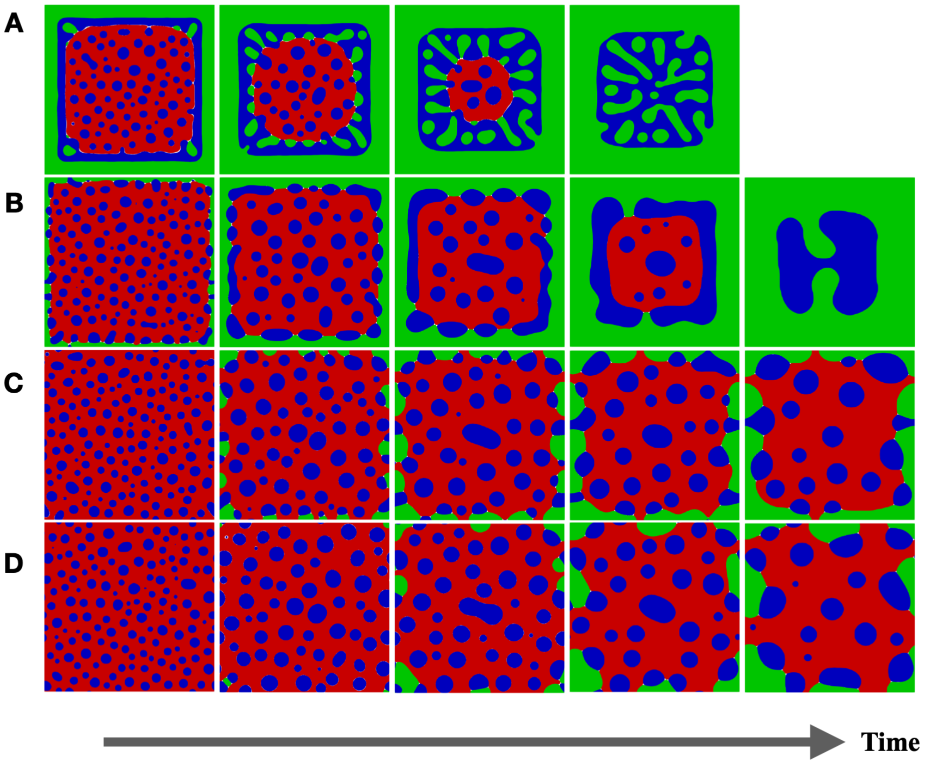


Figure 10: Evolution of phase-field variables $\varphi_i$ for **a slow dynamical regime** $\tau_d = 100\tau$ **with** $\tau_R = 100\tau$. From up to bottom: snapshots corresponding to simulation results with A) $\tau_P = \tau$; B) $\tau_P = 10\tau$; C) $\tau_P = 50\tau$; and D) $\tau_P = 100\tau$.

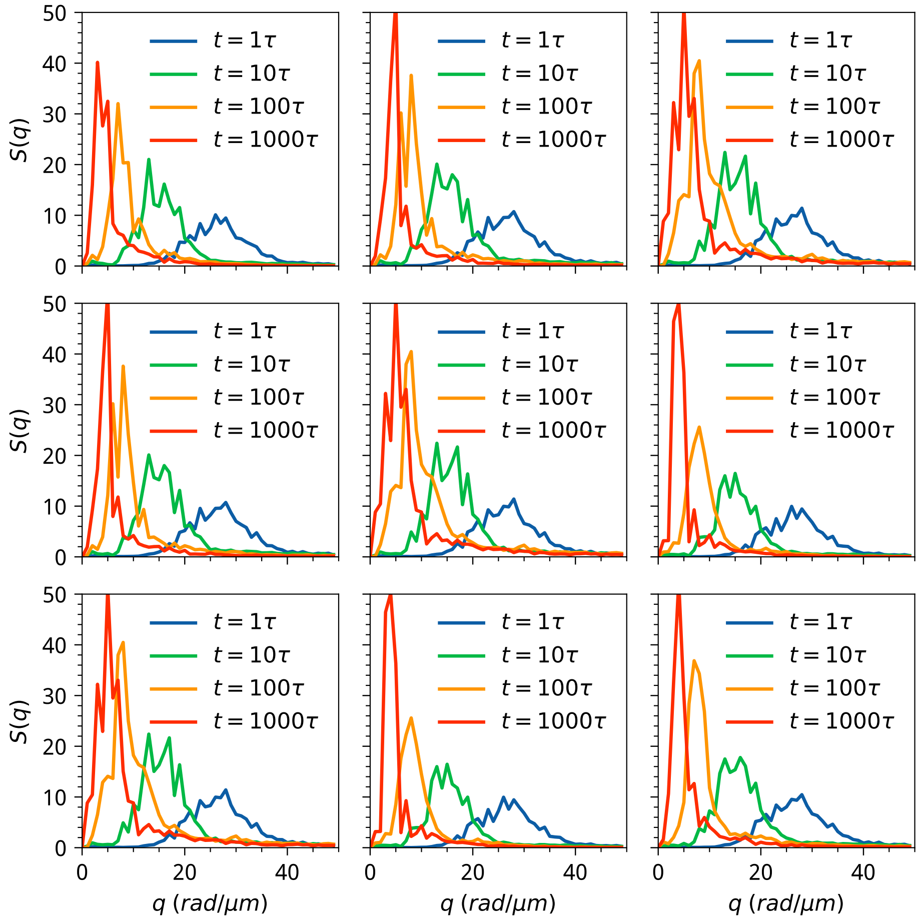


Figure 11: Structure factors for a fast dynamical regime $\tau_d = \tau$. Three panels corresponds to (A) Fixed transcription rate $\tau_R = 1$, varying translation from left to right: $\tau_P = \{1, 10, 100\}\ \tau$ (B) Fixed transcription rate $\tau_R = 10\tau$, varying translation from left to right: $\tau_P = \{1, 10, 100\}\ \tau$ (C) Fixed transcription rate $\tau_R = 100\tau$, varying translation from left to right: $\tau_P = \{1, 10, 100\}\ \tau$.

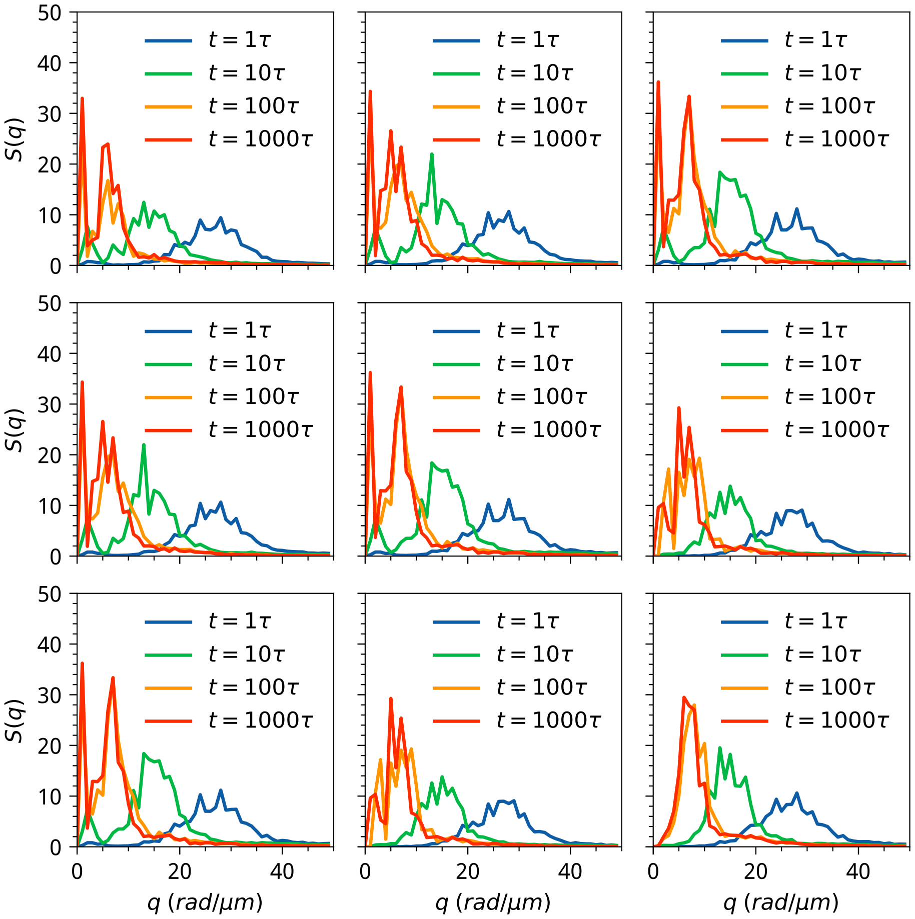


Figure 12: Structure factors for an intermediate dynamical regime $\tau_d = 10\tau$. Three panels corresponds to (A) Fixed transcription rate $\tau_R = \tau$, varying translation from left to right: $\tau_P = \{1, 10, 100\}\,\tau$ (B) Fixed transcription rate $\tau_R = 10\tau$, varying translation from left to right: $\tau_P = \{1, 10, 100\}\,\tau$ (C) Fixed transcription rate $\tau_R = 100\tau$, varying translation from left to right: $\tau_P = \{1, 10, 100\}\,\tau$.

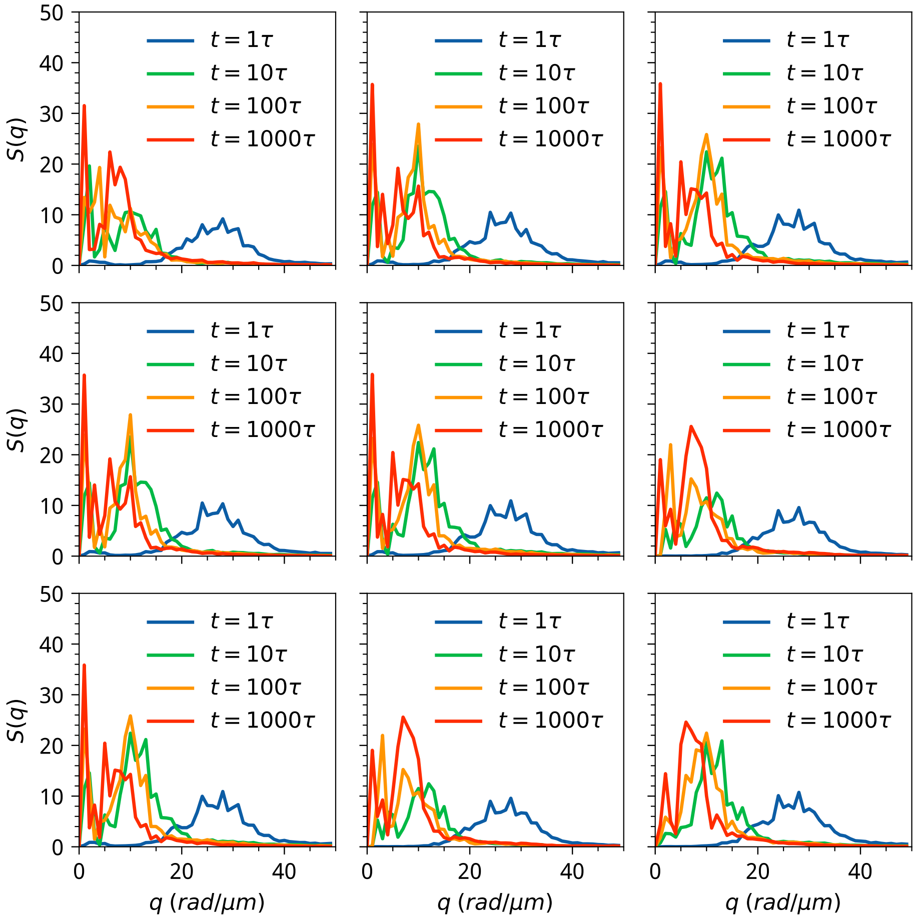


Figure 13: Structure factors for a slow dynamical regime $\tau_d = 100\tau$. Three panels corresponds to (A) Fixed transcription rate $\tau_R = \tau$, varying translation from left to right: $\tau_P = \{1, 10, 100\}\,\tau$ (B) Fixed transcription rate $\tau_R = 10\tau$, varying translation from left to right: $\tau_P = \{1, 10, 100\}\,\tau$ (C) Fixed transcription rate $\tau_R = 100\tau$, varying translation from left to right: $\tau_P = \{1, 10, 100\}\,\tau$.

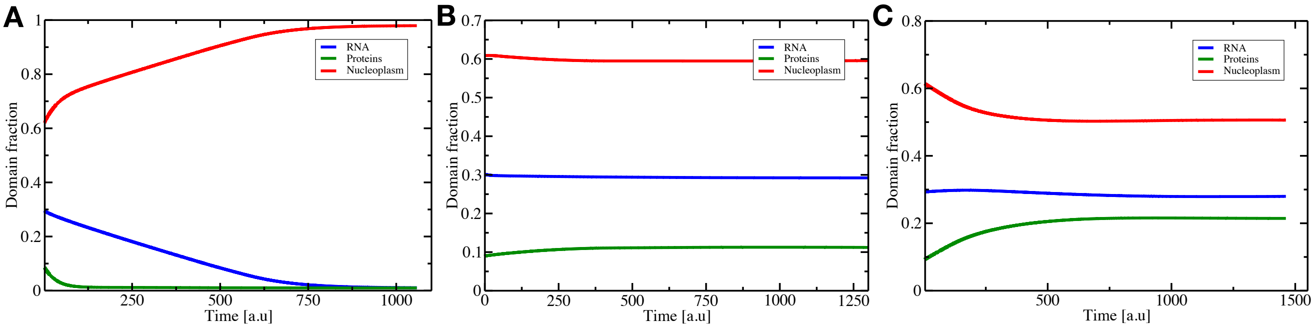


Figure 14: Temporal evolution of the average concentration of the three species $<\varphi_i>$ at different kinetic regimes with $\tau_R = \tau_P = 100\tau$: A) fast dynamical regime $\tau_d = \tau$; B) an intermediate dynamical regime $\tau_d = 10\tau$; C) slow dynamical regime $\tau_d = 100\tau$.
